# Unveiling Movement Intention after Stroke: Integrating EEG and EMG for Motor Rehabilitation

**DOI:** 10.1101/2024.02.22.581596

**Authors:** Eduardo López-Larraz, Andrea Sarasola-Sanz, Niels Birbaumer, Ander Ramos-Murguialday

**Author notes:** Correspondence: Eduardo López-Larraz.

## Abstract

Detecting attempted movements of a paralyzed limb is a key step for neural interfaces for motor rehabilitation and restoration after a stroke. In this paper, we present a systematic evaluation of electroencephalographic (EEG) and electromyographic (EMG) activity to decode when stroke patients with severe upper-limb paralysis attempt to move their affected arm. EEG and EMG recordings of 35 chronic stroke patients were analyzed. We trained classifiers to discriminate between rest and movement attempt states relying on brain, muscle, or both types of features combined. Our results reveal that: i) EEG and residual EMG features provide complementary information to detect attempted movements, obtaining significantly higher decoding accuracy when both sources of activity are combined; ii) EMG-based, but not EEG-based, decoding accuracy correlates with the degrees of impairment of the patient; and iii) the percentage of patients that achieve decoding accuracy above the chance level strongly depends on the type of features considered, and can be as low as 50% of them if only ipsilesional EEG is used. These results offer new perspectives to develop improved neurotechnologies that establish a more accurate contingent link between the central and peripheral nervous system after a stroke, leveraging Hebbian learning and facilitating functional plasticity and recovery.

## 1 Introduction

Neural interfaces have great potential for the rehabilitation of upper-limb paralysis after a stroke^1–4^. Many research laboratories worldwide are currently developing and testing body actuators controlled by brain and muscle activity^5–11^. Patients can use these systems to retrain their damaged motor network and improve their function. On the one hand, brain control of rehabilitative devices does not require any physical movement and can, therefore, be used even by patients with severe paralysis^12^. Furthermore, the synchronous activation of the brain regions responsible for movement and the stimulation of the limb with peripheral feedback can exploit plasticity mechanisms, eliciting motor recovery^13^. On the other hand, control strategies based on muscular activity can exploit the residual capabilities of each patient^14^, using the (sometimes minimal, but decodable) muscle contractions to actively drive rehabilitation^15–17^.

Studies that quantify the clinical efficacy of neurally controlled body actuators in stroke rehabilitation interventions are still scarce^2,18^. This is mainly due to the high amount of resources—in terms of number of patients, funding and time—required to conduct properly statistically-powered studies^19^. There are still numerous variables that need to be optimized to determine the most effective intervention for each patient^2^. For instance, it is unclear whether associative proprioceptive feedback dependent on ipsilesional activity leads to superior recovery compared to using contralesional or bihemispheric activity. Similarly, there is no experimental evidence of whether brain or muscle control of rehabilitative devices provides different recovery outcomes. Furthermore, there is currently no estimation of the percentage of patients who could satisfactorily benefit from this type of therapy, nor is there a successful triage strategy. The main reason for this is the lack of basic neuroscientific understanding of recovery after stroke in all phases (acute, sub-acute and chronic) and the potential mechanisms that could be leveraged to facilitate or induce motor recovery, especially in chronic patients.

For a rehabilitation scenario, where maximizing motor recovery is the goal, it is currently not possible to computationally simulate which type of training would be most effective, although some advances are being explored^20^. However, one can estimate offline the performance of algorithms that decode a patient’s movement intentions when relying on different sources of neural activity, determining the amount of neural control or contingency that could be obtained. Based on this information, future studies could adapt their experimental design according to the type of neural activity that seems to provide better information about the intended movement, to establish a better link (i.e., contingency) between central and peripheral neural activity, and facilitate Hebbian plasticity^5^.

In this study, we assessed the information provided by electroencephalography (EEG) and electromyography (EMG) to detect attempted hand movements in stroke patients, with the potential to use this information to control body actuators in an ecologically valid manner. We trained linear classifiers with different sets of features to discriminate between rest and movement attempt states and used them to answer three questions. First, we assessed if brain and muscle activity provide redundant or complementary information to detect the attempts of movement with the paralyzed hand. Second, we studied if the movement attempt decoding accuracy obtained relying on brain and/or muscle activity is related to the degrees of impairment of the patient. Third, we performed a more exhaustive analysis to compare 9 different combinations of features from EEG and/or EMG to quantify how many patients could control a neural interface based on these features and what their performance would be.

## 2 Materials and methods

### 2.1 Patients

Data from 35 chronic stroke patients with severe upper-limb paralysis (22 male, mean age 53.9±12.0 years, range 29–73, time since stroke 61.0±56.9 months, range 10–232) were analyzed in this study. These patients were recruited to participate in a clinical study investigating the rehabilitative effects of brain-machine interfaces (BMIs)^5^. From all the patients assessed for eligibility for the clinical trial (n=504), these 35 performed a screening assessment in which their EEG and EMG activity were recorded while they attempted to move their paralyzed hand in an open/close motion. The inclusion criteria were as follows: (1) hand paralysis with no finger extension; (2) minimum time since stroke of 10 months; (3) age between 18 and 80 years; (4) no psychiatric or neurological condition other than stroke; (5) no cerebellar lesion or bilateral motor deficit; (6) not pregnant; (7) no epilepsy or medication for epilepsy during the last 6 months; (8) eligibility to undergo magnetic resonance imaging (MRI); and (9) ability to understand and follow instructions. Further demographic and clinical data of the patients can be found in Table 1. The experiments were conducted at the University of Tübingen, Germany. The experimental procedure was approved by the ethics committee of the Faculty of Medicine of the University of Tübingen, and all the patients provided written informed consent.

**Table 1.**
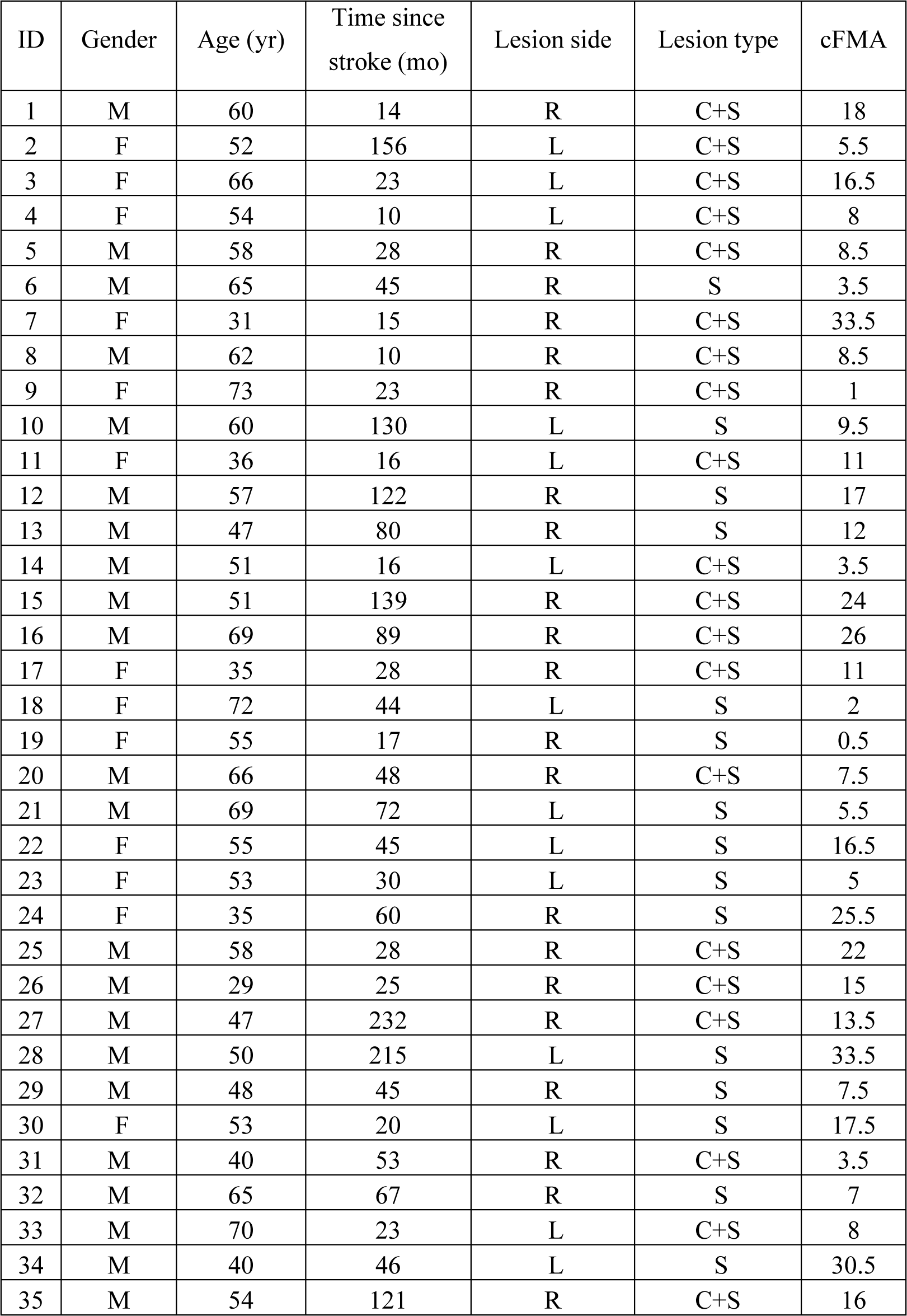
Demographic data. Lesion side indicates the affected brain hemisphere. Lesion type indicates if the stroke affected subcortical areas (S) or cortical and subcortical areas (C+S). cFMA stands for combined Fugl-Meyer assessment, comprising hand and arm motor scores combined, excluding coordination, speed and reflexes (range 0 - 54 points, with 54 points indicating normal hand/arm function).

### 2.2 Experimental design and procedure

Patients underwent one screening session in which their electroencephalographic (EEG) and electromyographic (EMG) activity were recorded while they attempted to open and close their completely paralyzed hand (Figure 1a). This screening session was conducted before the patients started the study described in Ramos-Murguialday et al., 2013^5^. Each patient completed between 4 and 6 blocks of 17 trials each. The number of blocks depended on the patients’ level of fatigue, as some of them requested to stop before finishing the planned 6 blocks. Audiovisual cues were displayed to guide the patients during the different phases of the trials (Figure 1b): rest (with a random duration between 4-5 seconds); movement attempt (4 seconds); and inter-trial interval (with random duration between 8-9 seconds). Patients were instructed to attempt to open and close their paretic hand at a comfortable pace during the movement attempt interval. They were informed about how to minimize activity that might contaminate EEG recordings (e.g., eye blinks, facial and jaw movements), as well as compensatory movements with the rest of the body.

**Figure 1.**
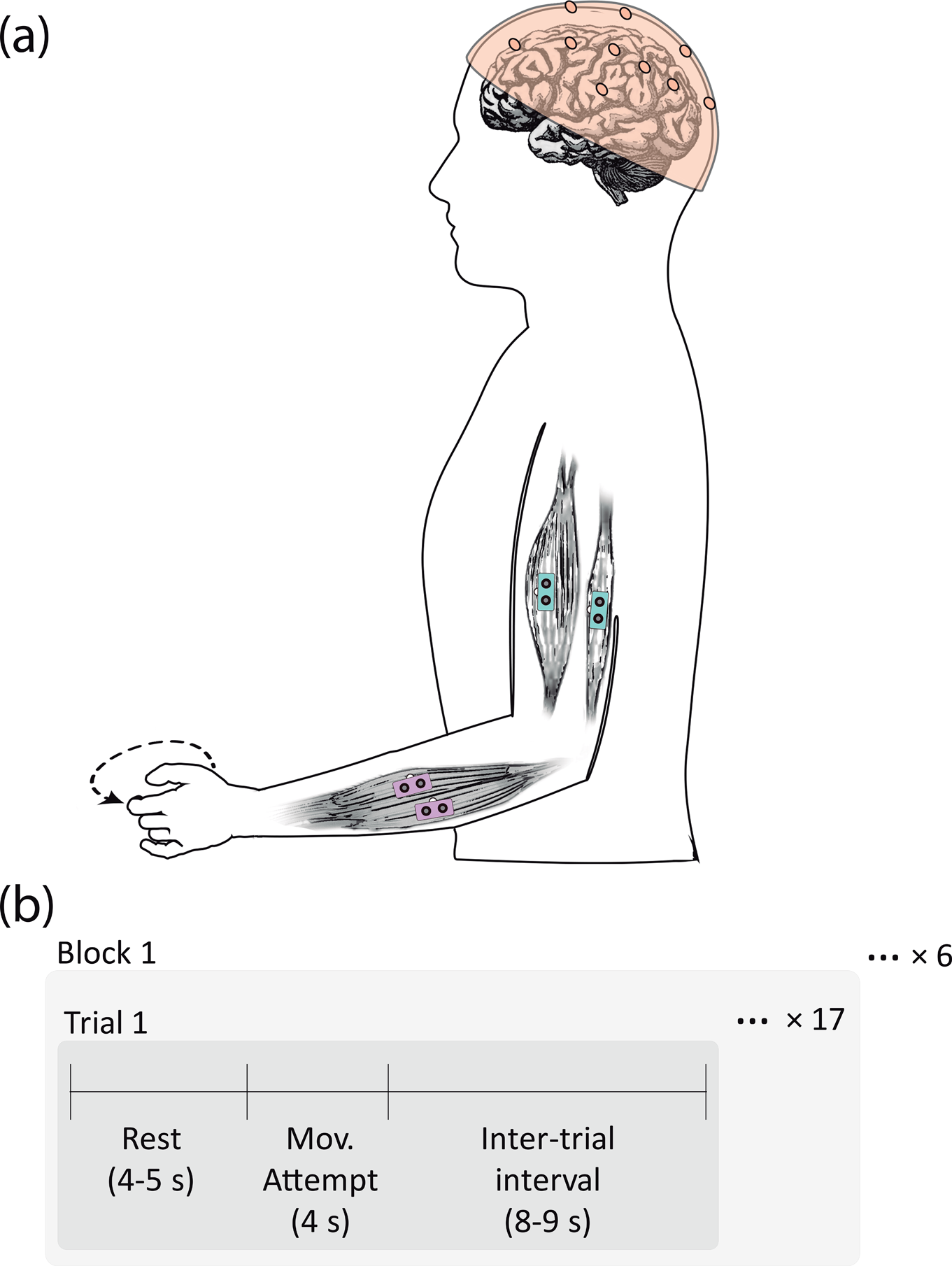
Experimental design. (a) Experimental setup, including the distribution of EEG and EMG electrodes used. (b) Time-diagram of the recording, describing the blocks, trials, and intervals within a trial. The audiovisual cues were provided at the beginning of each interval.

A commercial Acticap system (BrainProducts GmbH, Germany) was used to record EEG activity from 16 cortical positions (Fp1, Fp2, F3, Fz, F4, T7, C3, Cz, C4, T8, CP3, CP4, P3, Pz, P4, and Oz, according to the international 10/20 system), with the ground and reference placed on AFz and FCz, respectively. Vertical and horizontal electrooculography (EOG) was recorded to capture eye movements and for subsequent artifact removal. EMG activity was recorded with bipolar derivations from four muscle groups of each arm: *extensor carpi ulnaris*, *extensor digitorum*, external head of the *biceps* and external head of the *triceps*. All signals were synchronously recorded at 500 Hz using two BrainAmp amplifiers (BrainProducts GmbH, Germany).

### 2.3 Algorithm for detection of movement attempts

We implemented a procedure to decode when patients were attempting to move their paretic hand based on their EEG and/or EMG activity. By extracting features from various cortical positions and muscles and using them as inputs for a classifier, we aimed to characterize how the detection of movement attempts depends on the type of activity used.

In this analysis, we simulated a pseudo-online setup (i.e., it was tested offline but with all processing steps simulating an online scenario). We performed a block-based *N*-fold cross-validation separately for each patient, where *N* was the number of blocks for that patient (ranging from 4 to 6). For each fold, one block was separated as the test set, and the remaining blocks were used as the training set.

#### 2.3.1 Data preprocessing

The EEG signals were band-pass filtered between 0.1 and 48 Hz, while the EMG signals were high-pass filtered at 20 Hz (4-th order causal Butterworth filters). The EEG and EMG signals were then segmented into 7-second trials (from −3 to 4 seconds, with 0 being the presentation of the movement attempt audiovisual cue).

#### 2.3.2 Artifact removal

The trials in the training set underwent an artifact removal procedure to minimize the influence of contaminations in the data used to calibrate the classifier^21^. Ocular contaminations were removed from the EEG using a linear regression, as proposed by Schlögl et al^22^. Motion and muscle artifacts in the EEG were rejected using a threshold-based statistical procedure that discarded trials showing high power in delta or gamma frequencies (3 standard deviations above the mean) during either the rest or the movement attempt intervals. Similarly, trials with strong EMG activation during the rest interval or with EMG activations on the healthy arm (3 standard deviations above the mean) were discarded using a threshold-based statistical procedure. Further details of the algorithm for artifact removal can be found in López-Larraz et al., 2018^21^.

#### 2.3.3 Classifier training and evaluation

EEG and/or EMG features were extracted from one-second time windows to discriminate between the rest and movement attempt states. To train the classifier, we selected five windows from each trial of the training dataset to characterize the rest interval (i.e., interval [-2, 0] s with a sliding step of 0.25 s) and five windows for the movement attempt interval (i.e., interval [1, 3] s with a sliding step of 0.25 s). The movement attempt interval was defined as one second after the presentation of the cue to ensure that the patients had initiated the action, given their slow reaction time^23^.

EEG features were extracted from a set of electrodes (*S_EEG_*), which was defined for each of the different schemes presented in Section 2.4. The EEG signals were re-referenced with small Laplacian derivations^24^. The average alpha ([7-13] Hz) and beta ([14-30] Hz) power were calculated with an order-20 autoregressive model based on the Burg algorithm^25^, resulting in a total of [2×length(*S_EEG_*)] features for each window.

EMG features were extracted from a set of electrodes (*S_EMG_*), defined for each of the schemes presented in Section 2.4. EMG features were computed as the waveform length (WL) of the one-second windows from each electrode, resulting in [length(*S_EMG_*)] features per window.

The resulting feature vectors were z-score normalized, and the normalization parameters were stored and used to normalize the data in the test set. These feature vectors were then used to train a linear discriminant analysis (LDA) classifier.

The test trials were processed to remove ocular contaminations, using the parameters computed from the training trials (see Section 2.3.2). To simulate an online scenario, the test trials were evaluated with a one-second sliding window, from −3 to 4 seconds, applied every 20 ms (note that the first classifier output is generated at t = −2 s). The features of these one-second windows were extracted and normalized based on the parameters computed with the training data. The performance of the movement attempt classifier was measured in terms of average decoding accuracy. This average accuracy was computed as the mean of the correctly classified samples in the rest interval ([-2, 0] s) and in the movement attempt interval ([1, 4] s).

### 2.4 Data analysis and statistics

The results of movement attempt detection using different types of features were used to address three questions. First, whether brain and muscle activity provide redundant or complementary information for identifying attempted movements with the paralyzed hand. Second, whether the accuracy of decoding movement attempts obtained relying on brain and/or muscle activity correlates with the patient’s level of impairment. Third, we compared the number of patients who could control a neural interface for each of the 9 different combinations of EEG and/or EMG features, and what their performance would be.

#### 2.4.1 EEG and EMG to detect movement attempts: complementary or redundant?

In this initial analysis, we introduced three strategies for detecting movement attempts. We selected these three strategies as representative examples based on EEG, EMG, and a hybrid combination of EEG+EMG features. These strategies are rooted in the hypothesis that a neural interface capable of retraining the damaged motor circuits could enhance motor recovery through instrumental learning and Hebbian mechanisms.

The first strategy employs **ipsilesional EEG activity**, i.e., activity of the electrodes placed over the contralateral hemisphere to the hand that the patient is attempting to move (*S_EEG_* = [C3, CP3, P3] for patients with right hand paralysis, or [C4, CP4, P4] for patients with left hand paralysis; *S_EMG_* = []). This strategy has been previously proposed to train perilesional areas and attempt to reestablish the damaged circuits, strengthening their connections with the impaired limb^5,26^.

The second strategy utilizes the **EMG activity of the muscles that are “involved”** in the hand opening movement (*S_EEG_* = []; *S_EMG_* = [*extensor carpi ulnaris*, *extensor digitorum*]) to detect this action (we only placed electrodes on the extensors as they were the muscles that we wanted to reinforce, and because the hand was, by default, closed due to spasticity in almost all the cases, i.e., hyper-flexion). Movement-attempt decoding using EMG can be thought as a natural myoelectric control strategy for prostheses, relying on the residual activity of the muscles to detect the intended movement^14,27^.

The third strategy combined these two sources of activity (**ipsilesional EEG, and EMG of muscles “involved” in the movement**) to detect the movement attempts (*S_EEG_* = [C3, CP3, P3] for patients with right hand paralysis, or [C4, CP4, P4] for patients with left hand paralysis; *S_EMG_* = [*extensor carpi ulnaris*, *extensor digitorum*]). This way, it can take advantage of the potential complementary information from the brain and residual muscle activity to offer higher detection accuracy as well as to involve both central and peripheral structures to strengthen their link, as previously proposed in hybrid brain-machine interface studies^8^.

We examined the impact of the classifier’s input activity type on movement attempt detection by conducting a repeated measures analysis of variance (ANOVA). In this analysis, the decoding accuracy served as the dependent variable, while the within-subjects factor was the type of features employed (EEG, EMG, EEG+EMG). Greenhouse-Geisser correction was applied when the sphericity assumption was violated. Post-hoc paired t-tests were conducted with Bonferroni-Holm correction for multiple comparisons. The effect size of the significantly different pairs was computed using Cohen’s d. In addition, we computed Pearson’s correlation coefficient between each pair of decoding accuracy results to study potential significant relationships between the performance achieved using the different features.

#### 2.4.2 Relationship between decoding accuracy and motor impairment

Through this analysis, we explored whether the accuracy achievable in detecting movement attempts using EEG, EMG, or EEG+EMG corresponds to the level of a patient’s motor impairment. The degrees of impairment was quantified using the upper limb scores from the Fugl-Meyer assessment (excluding coordination, speed, and reflex-related scores to minimize variability^28^), as outlined in Table 1. For each of the three strategies presented, we calculated the Pearson’s correlation coefficient between decoding accuracy and the patients’ impairment level.

#### 2.4.3 Comparison of different strategies to detect the movement attempts

The three strategies previously described rely on features from ipsilesional EEG activity and EMG activity from the muscles involved in the task performed by the patients. However, other methodologies have also been employed in the literature to detect movement attempts, either for assistive or rehabilitative purposes. A comparison of performance across various feature sets for detecting movement attempts with a paralyzed limb can offer valuable insights for designing future neural interfaces. To conduct this analysis, we implemented and evaluated six additional strategies for detecting movement attempts:

##### Contralesional activity

Instead of employing ipsilesional electrodes, this approach utilizes the three electrodes over the contralesional hemisphere for feature extraction (*S_EEG_* = [C4, CP4, P4] for patients with right hand paralysis, or [C3, CP3, P3] for patients with left hand paralysis; *S_EMG_* = []).

##### Motor cortex activity

This approach combined EEG activity from the eight electrodes placed over both motor cortex hemispheres (*S_EEG_* = [C3, Cz, C4, CP3, CP4, P3, Pz, P4]; *S_EMG_*= []), regardless of stroke location.

##### Muscles unrelated to the task

In this strategy, we extracted features from the muscles that should not be activated during a hand opening-closing movement (*S_EEG_* = []; *S_EMG_* = [*biceps*, *triceps*]). This approach was assessed to explore whether compensatory muscle activation unrelated to the task contributes information for detecting movement attempts or introduces decoding bias.

##### All the recorded muscles

Here, features extracted from the four recorded muscles were combined (*S_EEG_* = []; *S_EMG_* = [*extensor carpi ulnaris*, *extensor digitorum, biceps*, *triceps*]) to assess how decoding accuracy changes when the activity of muscles related and unrelated to the task are combined.

##### Motor cortex activity + involved muscles

This approach combines the EEG activity of electrodes placed over the motor cortex with the muscles involved in the hand movement (*S_EEG_* = [C3, Cz, C4, CP3, CP4, P3, Pz, P4]; *S_EMG_* = [*extensor carpi ulnaris*, *extensor digitorum*]).

##### Motor cortex activity + all muscles

In this approach, the activity of the eight EEG electrodes over the motor cortex and the four EMG electrodes are combined to evaluate the performance of the strategy incorporating all features (*S_EEG_* = [C3, Cz, C4, CP3, CP4, P3, Pz, P4]; *S_EMG_* = [*extensor carpi ulnaris*, *extensor digitorum, biceps*, *triceps*]).

For all nine proposed strategies to detect movement attempts, we calculated three metrics: (1) the percentage of patients achieving decoding accuracy above chance level; (2) the average decoding accuracy for all patients; and (3) the average decoding accuracy for patients performing above chance level. This last metric provides an estimate of how effectively patients could control the neural interface, excluding those initially considered as non-responders^29^.

## 3 Results

The data from one patient (P22) was excluded due to excessive artifacts. Consequently, all subsequent analyses are based on the data of 34 patients.

### 3.1 EEG and EMG provide complementary information to detect movement attempts

The decoding accuracy varied significantly depending on the type of features utilized by the classifier, as revealed by the repeated measures ANOVA (*F*(1.19, 39.22) = 7.42; *p* = 0.007). The average temporal response of the three basic strategies for detecting the movement attempts—based on EEG (ipsilesional activity), EMG (involved muscles) and EEG+EMG (ipsilesional + involved muscles)—is depicted in Figure 2a, while the mean decoding accuracy of each approach (averaging performance during rest and movement attempt intervals) is displayed in Figure 2b. Post-hoc paired comparisons evidenced significantly different decoding accuracy between detection based on EEG and EEG+EMG features (*t*(33) = −4.7, *p* = 0.0001; Cohen’s *d* = 0.71), as well as between detection based on EMG and EEG+EMG features (*t*(33) = −2.5, *p* = 0.037; Cohen’s *d* = 0.28).

**Figure 2.**
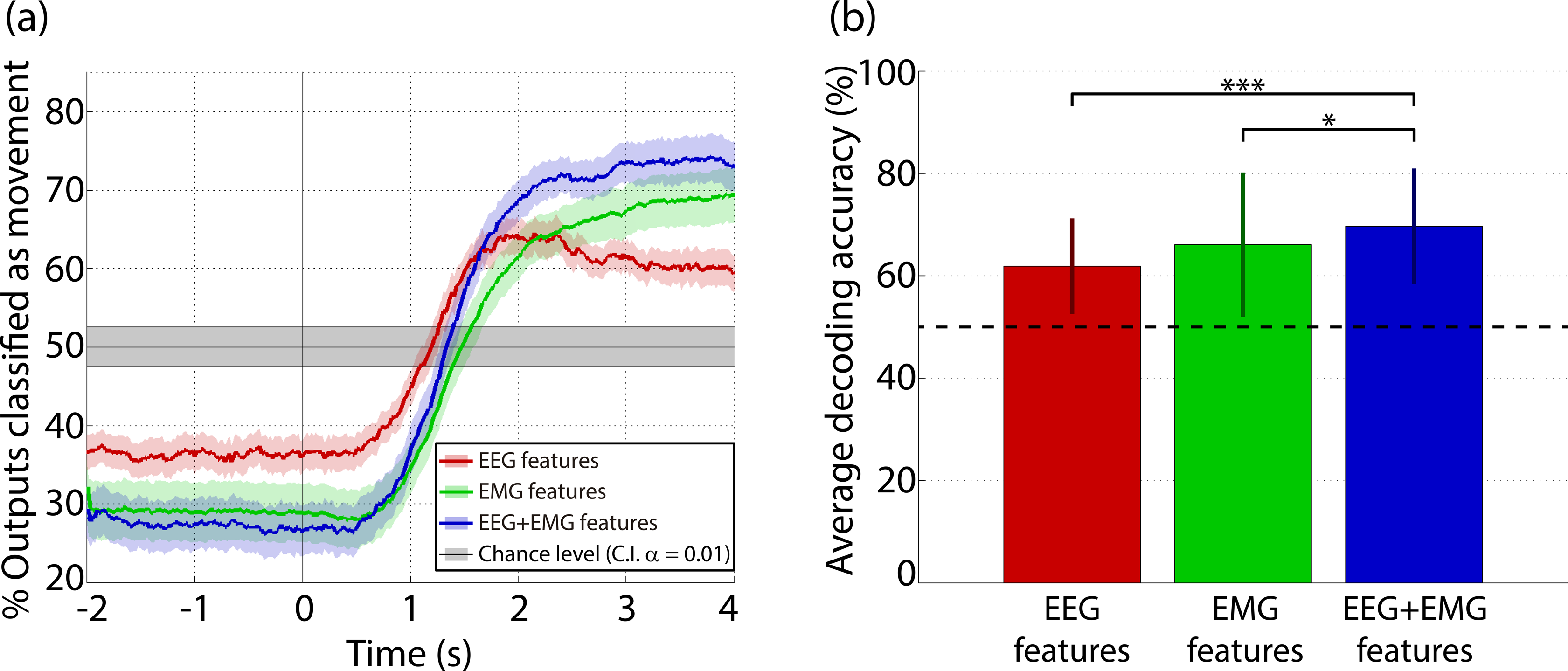
Decoding accuracy for the three basic schemes: ipsilesional EEG, EMG of involved muscles and the combination of ipsilesional EEG and EMG of involved muscles. (a) Temporal response of the classifiers trained with EEG, EMG, or EEG+EMG features. The lines illustrate the percentage of classifier outputs labeled as movement, averaged across all patients, while the shades indicate the standard error of the mean. Values before t = 0 correspond to false positives, while values after t = 0 correspond to true positives. The shaded gray area indicates the confidence interval of the chance level (alpha = 0.01), computed on the basis of all test trials^30^. (b) Accuracy for each feature configuration, averaged across rest ([-2, 0] s) and movement attempt ([1, 4] s) intervals. The vertical bars denote the standard deviation. Significantly different pairs of results are marked: *: p<0.05; ***: p<0.001.

The performance of the classifier trained using EEG+EMG features correlated significantly with the classifier trained using EEG features (*r* = 0.58, *p* = 0.0003) and with the classifier trained using EMG features (*r* = 0.80, *p* = 1.53×10^-8^). However, the performance achieved with EEG and with EMG features were not significantly correlated (*r* = 0.12, *p* = 0.50), suggesting that EEG and EMG features provide complementary information. Figure 3 illustrates the accuracy achieved with each of the three decoding strategies for each patient, revealing the relationship between high accuracies using EEG/EMG features (*x*-axis/*y*-axis, respectively) and using the combination of EEG+EMG features (color-coded).

**Figure 3.**
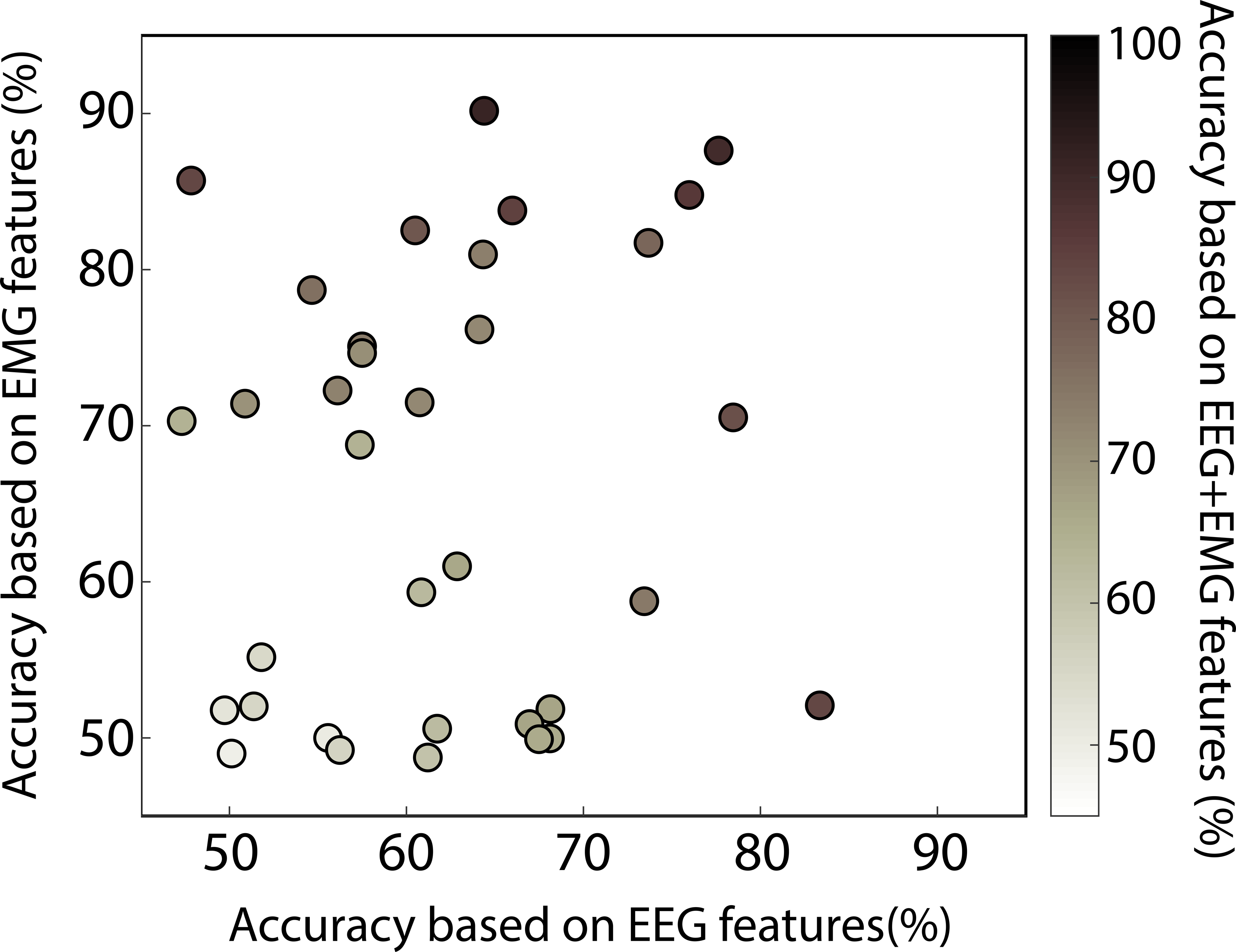
Relationship between the accuracy achieved with the three types of features. Distribution of decoding accuracy values for each patient using EEG features (x-axis), EMG features (y-axis) or EEG+EMG features (color-coded).

### 3.2 EMG-based but not EEG-based decoding accuracy correlates with motor impairment

We investigated whether the performance obtained with each type of feature (ipsilesional EEG, EMG of involved muscles, and their combination) is related to the patient’s impairment level. Figure 4 presents the results of the correlation analysis between the performance of the classifier for each feature configuration and the motor impairment, as measured by the Fugl-Meyer assessment. The degrees of motor impairment of the patients significantly correlated with their achieved performance using EMG features and EEG+EMG features (*r* = 0.51, *p* = 0.002 for EMG, Figure 4b; *r* = 0.39, *p* = 0.023 for EEG+EMG, Figure 4c), but not when using EEG features (*r* = 0.09, *p* = 0.61, Figure 4a).

**Figure 4.**
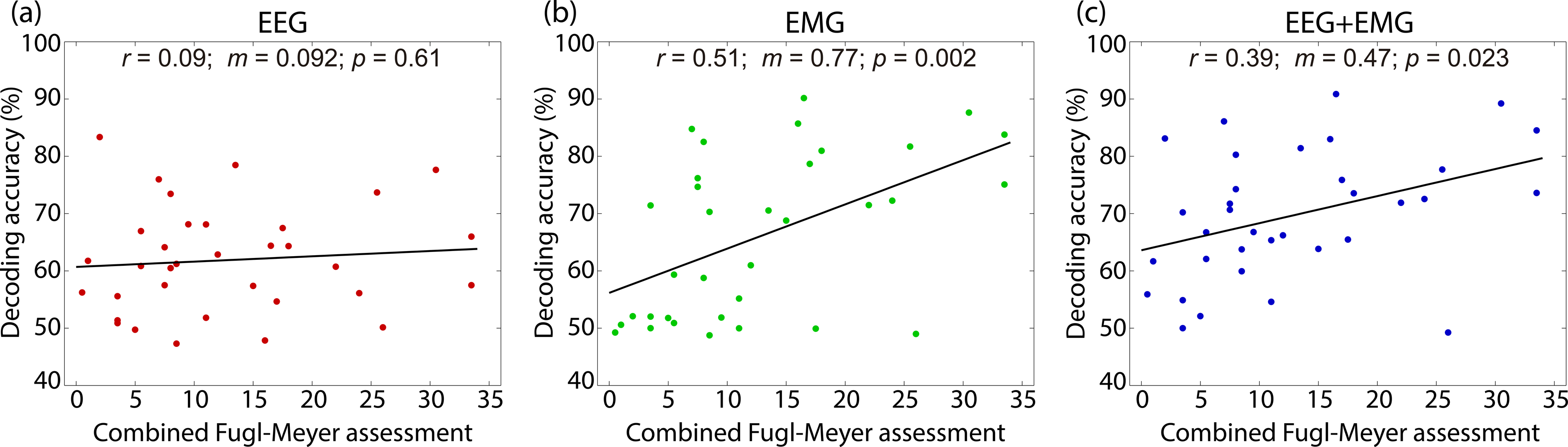
Correlation between decoding accuracy and motor impairment. (a) Correlation of decoding accuracy achieved with EEG features and cFMA score. (b) Correlation of decoding accuracy achieved with EMG features and cFMA score. (c) Correlation of decoding accuracy achieved with EEG+EMG features and cFMA score. Each dot corresponds to one patient. The correlation value (r), the slope of the least-squares regression line of best fit (m) and the p-value of the correlation (p) are displayed at the top of each panel.

### 3.3 Differences in usability and performance for the different types of features

Figure 5 provides a comprehensive summary of all feature combinations and their respective outcomes. Initially, we assessed the number of patients achieving performance significantly above the chance level. The confidence interval of the chance level was computed individually for each patient depending on their number of recorded trials^30^. When utilizing ipsilesional EEG activity alone, only 50% of patients achieved performance beyond chance, whereas 53% achieved this with EMG activity from the involved muscles. However, combining both these features raised the number of patients achieving significant control to 79% (columns 1, 4 and 7, in the second row of Figure 5). The percentage of users performing above chance increased further with the inclusion of bilateral EEG electrodes or when considering activity from muscles unrelated to the task.

**Figure 5.**
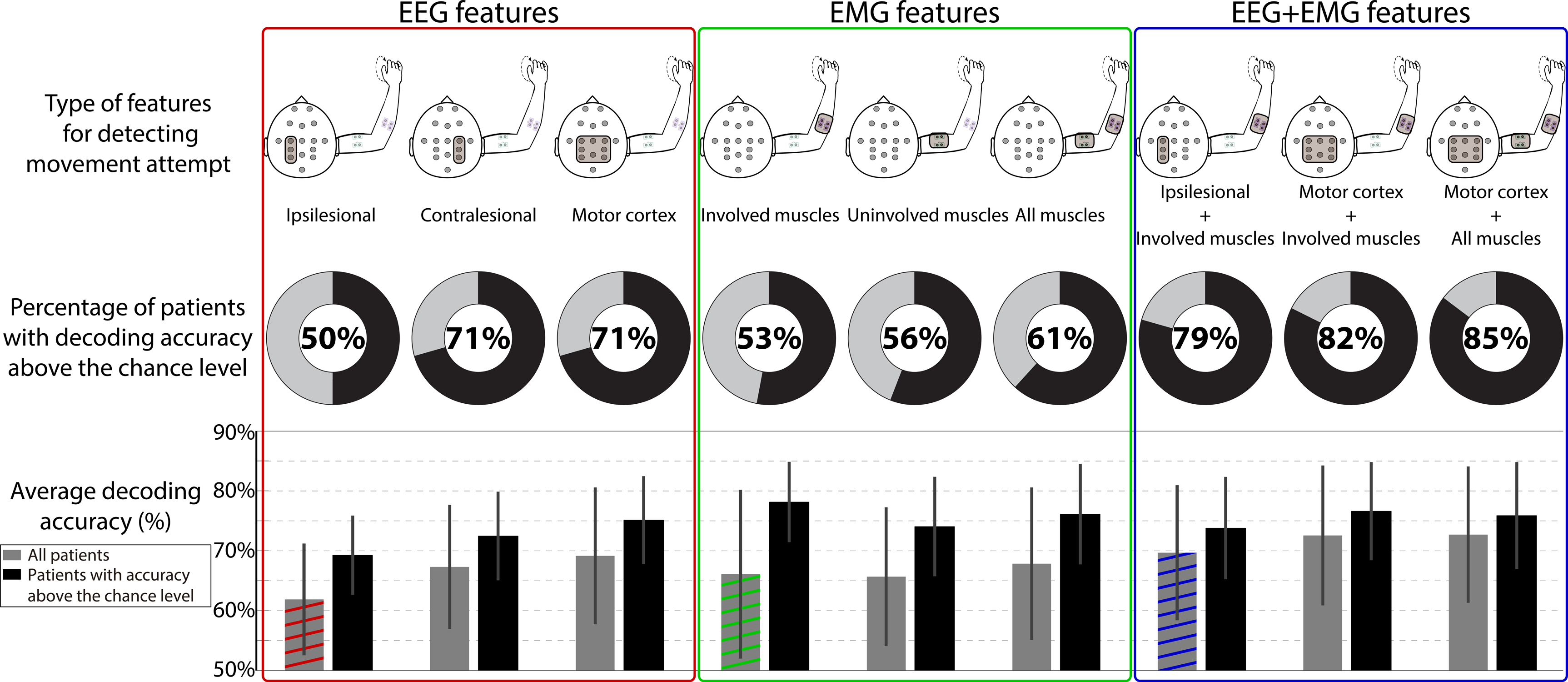
Detailed decoding accuracy results for all the feature combinations. The first row presents the 9 feature combinations used for detecting movement attempts. The second row indicates the percentage of patients surpassing chance-level accuracy. The third row displays the average decoding accuracy (mean and standard deviation) for each feature combination. Gray bars represent all patients, while black bars correspond to patients performing above chance. Red/green/blue-colored bars correspond to the same result as displayed in Figure 2.

The last row in Figure 5 details the average decoding accuracy for each feature combination. In terms of EEG features, accuracy using contralesional or bilateral features was 5.4% and 7.3% higher, respectively, than using ipsilesional features. Conversely, using EMG features from muscles related or unrelated to the task (or all combined) yielded nearly equivalent performance. As shown in section 3.1, the strategies that combined EEG and EMG features resulted in higher decoding accuracies.

As expected, excluding the patients whose performance was not significantly different to chance level led to an increase in performance, which ranged from 3.2% for the strategy utilizing all available features, (rightmost bars at the bottom of Figure 5) to 12.1% for the strategy based on EMG features of the involved muscles (leftmost bars in the green panel of Figure 5).

## 4 Discussion

This paper presented the first systematic evaluation of EEG and EMG activity for detecting movement attempts of a paralyzed limb in chronic stroke patients. In light of the increasing interest in neural interfaces for motor rehabilitation and assistive device control, these findings introduce new perspectives to improve future developments of these technologies.

Firstly, our results demonstrate the complementarity of EEG and residual EMG activity in detecting movement attempts of stroke patients, aligning with previous research. Balasubramanian and colleagues reported a poor temporal agreement between EMG-based and EEG-based movement decoding during brain-controlled robotic exoskeleton training, suggesting that these approaches may be detecting different processes^31^. In Spüler et al., 2018, we conducted a comparative analysis of methods for movement attempt decoding using EEG or EMG in chronic stroke patients during BMI robotic orthosis control ^32^. The results of that study showed that the best EEG decoder outperformed the best EMG decoder. In the study presented here, we analyzed data during movement attempts without the presence of assisted robotic movement, thus eliminating proprioceptive and haptic feedback (robotic movement of the hand), which could influence afference-related brain oscillatory activity and might explain the differences with the results presented in Spüler et al. The combination of EEG and EMG activity has been demonstrated to outperform movement detection based solely on EEG in studies with healthy subjects^33^. In a preliminary study, we also observed this result in a subset of patients from this study who had cortical stroke only^34^, representing the population of patients with the lowest BMI performance^35^. Taken together, these findings justify the further development of hybrid cortico-muscular neural interfaces, as they have the potential to better detect patients’ intentions. Various methods for combining EEG and EMG have been proposed: including the concatenation of EEG and EMG features (as proposed in this and other studies^33,34^), EEG-gated control based on EMG decoding of different degreess of freedom^8^, or EMG-gated control based on EEG classification of start-stop states^36^. Further research should elucidate the optimal approach for combining these signals to maximize system performance and rehabilitative efficacy.

The correlation analysis with clinical variables revealed that movement decoding based on EMG activity of the paralyzed muscles correlates significantly with impairment, while EEG decoding based on ipsilesional activity does not^a^. Previous studies have also reported differences in EMG patterns (and in EMG-based movement decoding accuracy) dependent on the motor impairment of stroke patients^14,37,38^, which makes sense considering that EMG is the behavioral output of the sensorimotor nervous system. In contrast, the analyzed EEG patterns and the decoding accuracy achieved with them seem to be primarily influenced by other factors, such as stroke location^35,39,40^, and may not directly reflect the motor impairment itself but could indicate neural impairment. Nevertheless, recent studies have highlighted the relevance of EEG oscillations as biomarkers of recovery^41,42^, suggesting an interplay between EEG sensorimotor modulation (within, and balance towards the ipsilesional hemisphere), EMG activity and clinical progress.

A relevant finding was that only 50% of the patients achieved performance above chance level using their ipsilesional EEG activity, and only 53% did so using EMG activity from their involved muscles. However, when these two types of features were combined, the number of patients achieving significant control increased to 79%. Thus, the combination of both brain and muscle activity not only enhances decoding accuracy but also expands the number of patients who would achieve accurate control through the neural interface. This could lead to greater effectiveness in the use of BMIs for stroke rehabilitation.

A common point of contention in BMI studies for stroke rehabilitation is whether feedback should be based on activity from the ipsilesional hemisphere^5,26^, the contralesional hemisphere^7^, or a combination of both^6,43,44^. Our results indicate that ipsilesional activity provides lower decoding accuracy, and fewer patients achieve performance above chance using this approach. Motor recovery following a stroke has been linked to increased ipsilesional activity and decreased contralesional activity^41,45,46^, which has motivated BMI approaches focused on strengthening ipsilesional activation^5,26^. Methods based on detecting movement attempts using contralesional (i.e., ipsilateral) activity may not directly address the interhemispheric imbalance caused by the stroke^47,48^. However, these approaches have shown positive results in terms of recovery, possibly due to intensive therapy, a common confounding factor in studies lacking a proper control group.

The study by Ramos-Murguialday et al., 2013, demonstrated that after ipsilesional BMI training, bihemispheric changes in function and structure occur in chronic stroke patients^5,49^. Therefore, the bihemispheric interplay and its involvement in recovery is clear. Our results do not shed light on which approach yields superior recovery, but provide valuable insights into the reliability of ipsilesional or contralesional EEG for detecting movement attempts. Based on these results, we speculate that combined strategies aiming to reinforce ipsilesional activity, but that rely on contralesional (or combined ipsi- and contralesional) activity when ipsilesional activity is insufficient, may be the most promising approach for brain-machine interfaces. This approach could benefit a larger number of patients by establishing a more reliable contingent connection between the brain, muscles, and movement. However, further investigation is needed to fully assess the rehabilitative potential of each approach.

The use of information from the muscles that should be involved in the attempted movement yielded similar results (avg. accuracy = 66.1%; percentage of patients above the chance level = 53%) when compared to ipsilesional EEG activity (avg. accuracy = 61.9%; percentage of patients above the chance level = 50%). Remarkably, even the muscles that should not be involved in this movement produced comparable results (avg. accuracy = 65.7%; percentage of patients above the chance level = 56%). This last finding reveals strong compensatory activity when patients attempt to perform a movement with their paralyzed limb. In principle, EMG biofeedback approaches should not rely on the activation of these unrelated muscles to control rehabilitative devices, as they might reinforce maladaptive plasticity and pathological synergies^50,51^. Nevertheless, monitoring this activity can be leveraged to provide negative feedback, teaching patients to reduce these compensatory mechanisms and normalize their abnormal muscle activation patterns^52,53^, for instance, by employing encoders trained on a healthy muscle activity pattern^27,54^.

In our analyses, we presented decoding accuracy as a means to quantify the informativeness of each source of activity in detecting movement attempts. Nonetheless, we cannot assume a direct causality between accuracy and the effectiveness of an approach for rehabilitation. Recent evidence has suggested a relationship between EEG-based decoding accuracy and functional recovery during a BMI intervention for stroke patients^6^. However, this only implies that for a fixed decoding strategy as the one proposed by Biasiucci et al.^6^, better control resulted in greater improvements. It remains to be determined whether neural interfaces relying on different sources of activity—which as we proved, would provide different performance—would have comparable influence in terms of rehabilitative potential. This question will need to be addressed in future meta-analyses, which should also consider other critical factors such as duration and intensity^55^. Additionally, these analyses could benefit from insights provided by new computational recovery models^20^.

Given recent evidence suggesting limitations in robotic rehabilitation compared to usual care for patients with severe paralysis^56^, it is crucial for this field to explore new directions that enhance the prospects of successful rehabilitation and improve efficacy. Neural interfaces offer the potential to leverage plasticity mechanisms^13^ and instrumental learning^57^ to facilitate recovery, serving as valuable neuroscientific tools for studying post-stroke learning and recovery. The integration of physiological measurements with rehabilitative devices holds significant promise for validation in clinical trials and eventual adoption as a recognized treatment for stroke recovery. This study has contributed valuable insights into how different measures of brain and muscle activity can be harnessed to detect movement intentions. Such insights may prove instrumental in the development of future therapies for stroke rehabilitation and the control of assistive body actuators based on neural interfaces.

## Acknowledgments

The authors thank all the people involved in the data recording for their hard work. This study was funded by the *f*ortüne-Program of the University of Tübingen (2422-0-1), the Eurostars Project E! 113928 SubliminalHomeRehab (SHR), the Bundesministerium für Bildung und Forschung (BMBF) (FKZ: SHR 01QE2023; REHOME 16SV8606 and REHALITY 13GW0213) and the European Union H2020-FETPROACT-EIC-2018-2020 (MAIA 951910).

## Conflicts of interest

Author ELL is employed by the company Bitbrain.

## Footnotes

a Despite not described in the results section, the accuracy achieved using only contralesional EEG electrodes, or all the motor cortex electrodes did not correlate with impairment either.

## References

1. Wang, W. et al. Neural Interface Technology for Rehabilitation: Exploiting and Promoting Neuroplasticity. Phys. Med. Rehabil. Clin. N. Am. 21, 157–178 (2010).

2. López-Larraz, E., Sarasola-Sanz, A., Irastorza-Landa, N., Birbaumer, N. & Ramos-Murguialday, A. Brain-machine interfaces for rehabilitation in stroke: a review. NeuroRehabilitation 43, 77–97 (2018).

3. Lin, D. J., Finklestein, S. P. & Cramer, S. C. New Directions in Treatments Targeting Stroke Recovery. Stroke 49, 3107–3114 (2018).

4. Vidaurre, C. et al. Challenges of neural interfaces for stroke motor rehabilitation. Front. Hum. Neurosci. 17, 1070404 (2023).

5. Ramos-Murguialday, A. et al. Brain-machine interface in chronic stroke rehabilitation: a controlled study. Ann. Neurol. 74, 100–108 (2013).

6. Biasiucci, A. et al. Brain-actuated functional electrical stimulation elicits lasting arm motor recovery after stroke. Nat. Commun. 9, 2421 (2018).

7. Bundy, D. T. et al. Contralesional Brain-Computer Interface Control of a Powered Exoskeleton for Motor Recovery in Chronic Stroke Survivors. Stroke 48, 1908–1915 (2017).

8. Sarasola-Sanz, A. et al. A Hybrid Brain-Machine Interface based on EEG and EMG activity for the Motor Rehabilitation of Stroke Patients. in 15th International Conference on Rehabilitation Robotics (ICORR) 895–900 (2017). doi:10.1109/ICORR.2017.8009362.

9. Dipietro, L. et al. Customized interactive robotic treatment for stroke: EMG-triggered therapy. IEEE Trans. Neural Syst. Rehabil. Eng. 13, 325–334 (2005).

10. de Kroon, J. R. & IJzerman, M. J. Electrical stimulation of the upper extremity in stroke: cyclic versus EMG-triggered stimulation. Clin. Rehabil. 22, 690–697 (2008).

11. Qian, Q. et al. Early stroke rehabilitation of the upper limb assisted with an electromyography-driven neuromuscular electrical stimulation-robotic arm. Front. Neurol. 8, 447 (2017).

12. Birbaumer, N., Ramos-Murguialday, A. & Cohen, L. Brain-computer interface in paralysis. Curr. Opin. Neurol. 21, 634–638 (2008).

13. Jackson, A. & Zimmermann, J. B. Neural interfaces for the brain and spinal cord— restoring motor function. Nat. Rev. Neurol. 8, 690–699 (2012).

14. Ramos-Murguialday, A. et al. Decoding upper limb residual muscle activity in severe chronic stroke. Ann. Clin. Transl. Neurol. 2, 1–11 (2015).

15. Sarasola-Sanz, A. et al. EMG-based multi-joint kinematics decoding for robot-aided rehabilitation therapies. in 14th International Conference on Rehabilitation Robotics 229–234 (2015). doi:10.1109/ICORR.2015.7281204.

16. Cesqui, B., Tropea, P., Micera, S. & Krebs, H. EMG-based pattern recognition approach in post stroke robot-aided rehabilitation: a feasibility study. J. Neuroeng. Rehabil. 10, 75 (2013).

17. Ho, N. S. K. et al. An EMG-driven exoskeleton hand robotic training device on chronic stroke subjects: Task training system for stroke rehabilitation. in 12th International Conference on Rehabilitation Robotics (ICORR) (2011). doi:10.1109/ICORR.2011.5975340.

18. Cervera, M. A. et al. Brain-computer interfaces for post-stroke motor rehabilitation: a meta-analysis. Ann. Clin. Transl. Neurol. 5, 651–663 (2018).

19. Chaudhary, U., Birbaumer, N. & Ramos-Murguialday, A. Brain-computer interfaces for communication and rehabilitation. Nat. Rev. Neurol. 12, 513–525 (2016).

20. Norman, S. L., Wolpaw, J. R. & Reinkensmeyer, D. J. Targeting neuroplasticity to improve motor recovery after stroke: an artificial neural network model. Brain Commun. 4, fcac264 (2022).

21. López-Larraz, E. et al. Event-related desynchronization during movement attempt and execution in severely paralyzed stroke patients: an artifact removal relevance analysis. NeuroImage Clin. 20, 972–986 (2018).

22. Schlögl, A. et al. A fully automated correction method of EOG artifacts in EEG recordings. Clin. Neurophysiol. 118, 98–104 (2007).

23. Yilmaz, O., Birbaumer, N. & Ramos-Murguialday, A. Movement related slow cortical potentials in severely paralyzed chronic stroke patients. Front. Hum. Neurosci. 8, 1033 (2015).

24. Hjorth, B. An on-line transformation of EEG scalp potentials into orthogonal source derivations. Electroencephalogr. Clin. Neurophysiol. 39, 526–530 (1975).

25. Burg, J. P. Maximum Entropy Spectral Analysis. in Proceedings of the 37th Annual International SEG Meeting (1975).

26. Pichiorri, F. et al. Brain-computer interface boosts motor imagery practice during stroke recovery. Ann. Neurol. 77, 851–865 (2015).

27. Sarasola-Sanz, A. et al. Design and effectiveness evaluation of mirror myoelectric interfaces: a novel method to restore movement in hemiplegic patients. Sci. Rep. 8, 16688 (2018).

28. Crow, J. L. & Harmeling-Van Der Wel, B. C. Hierarchical Properties of the Motor Function Sections of the Fugl-Meyer Assessment Scale for People After Stroke: A Retrospective Study Background and Purpose. The upper-extremity (UE) and lower-extremity. Phys. Ther. 88, 1554–1567 (2008).

29. Vidaurre, C. & Blankertz, B. Towards a cure for BCI illiteracy. Brain Topogr. 23, 194– 198 (2010).

30. Müller-Putz, G. R., Scherer, R., Brunner, C., Leeb, R. & Pfurtscheller, G. Better than random? A closer look on BCI results. Int. J. Bioelectromagn. 10, 52–55 (2008).

31. Balasubramanian, S., Garcia-Cossio, E., Birbaumer, N., Burdet, E. & Ramos-Murguialday, A. Is EMG a viable alternative to BCI for detecting movement intention in severe stroke? IEEE Trans. Biomed. Eng. 65, 2790–2797 (2018).

32. Spüler, M., López-Larraz, E. & Ramos-Murguialday, A. On the design of EEG-based movement decoders for completely paralyzed stroke patients. J. Neuroeng. Rehabil. 15, 110 (2018).

33. Leeb, R., Sagha, H., Chavarriaga, R. & Millán, J. del R. A hybrid brain-computer interface based on the fusion of electroencephalographic and electromyographic activities. J. Neural Eng. 8, 025011 (2011).

34. López-Larraz, E., Birbaumer, N. & Ramos-Murguialday, A. A hybrid EEG-EMG BMI improves the detection of movement intention in cortical stroke patients with complete hand paralysis. in 40th Annual International Conference of the IEEE Engineering in Medicine and Biology Society (EMBC) 2000–2003 (2018). doi:10.1109/EMBC.2018.8512711.

35. López-Larraz, E. et al. Stroke lesion location influences the decoding of movement intention from EEG. in 39th Annual International Conference of the IEEE Engineering in Medicine and Biology Society (EMBC) 3065–3068 (2017). doi:10.1109/EMBC.2017.8037504.

36. Bhagat, N. A. et al. Design and optimization of an EEG-based brain machine interface (BMI) to an upper-limb exoskeleton for stroke survivors. Front. Neurosci. 10, 122 (2016).

37. Zhang, X. & Zhou, P. High-density myoelectric pattern recognition toward improved stroke rehabilitation. IEEE Trans. Biomed. Eng. 59, 1649–1657 (2012).

38. Lee, S. W., Wilson, K. M., Lock, B. A. & Kamper, D. G. Subject-specific myoelectric pattern classification of functional hand movements for stroke survivors. IEEE Trans. Neural Syst. Rehabil. Eng. 19, 558–566 (2011).

39. Park, W., Kwon, G. H., Kim, Y.-H., Lee, J.-H. & Kim, L. EEG response varies with lesion location in patients with chronic stroke. J. Neuroeng. Rehabil. 13, 21 (2016).

40. López-Larraz, E., Ray, A. M., Birbaumer, N. & Ramos-Murguialday, A. Sensorimotor rhythm modulation depends on resting-state oscillations and cortex integrity in severely paralyzed stroke patients. in 9th International IEEE EMBS Conference on Neural Engineering (NER) 37–40 (2019). doi:10.1109/NER.2019.8717112.

41. Ray, A. M., Figueiredo, T. C., López-Larraz, E., Birbaumer, N. & Ramos-Murguialday, A. Brain oscillatory activity as a biomarker of motor recovery in chronic stroke. Hum. Brain Mapp. 41, 1296–1308 (2020).

42. Stinear, C. M. Prediction of motor recovery after stroke: advances in biomarkers. Lancet Neurol. 16, 826–836 (2017).

43. Ang, K. K. et al. A Randomized Controlled Trial of EEG-Based Motor Imagery Brain-Computer Interface Robotic Rehabilitation for Stroke. Clin. EEG Neurosci. 46, 310–320 (2015).

44. Ono, T. et al. Brain-computer interface with somatosensory feedback improves functional recovery from severe hemiplegia due to chronic stroke. Front. Neuroeng. 7, 19 (2014).

45. Ward, N. S., Brown, M. M., Thompson, A. J. & Frackowiak, R. S. J. Neural correlates of outcome after stroke: A cross-sectional fMRI study. Brain 126, 1430–1448 (2003).

46. Ward, N. S., Brown, M. M., Thompson, A. J. & Frackowiak, R. S. J. Neural correlates of motor recovery after stroke: a longitudinal fMRI study. Brain 126, 2476–2496 (2003).

47. Dodd, K. C., Nair, V. A. & Prabhakaran, V. Role of the Contralesional vs. Ipsilesional Hemisphere in Stroke Recovery. Front. Hum. Neurosci. 11, 469 (2017).

48. Murase, N., Duque, J., Mazzocchio, R. & Cohen, L. G. Influence of interhemispheric interactions on motor function in chronic stroke. Ann. Neurol. 55, 400–409 (2004).

49. Caria, A. et al. Brain–Machine Interface Induced Morpho-Functional Remodeling of the Neural Motor System in Severe Chronic Stroke. Neurotherapeutics (2019) doi:10.1007/s13311-019-00816-2.

50. García-Cossio, E., Broetz, D., Birbaumer, N. & Ramos-Murguialday, A. Cortex integrity relevance in muscle synergies in severe chronic stroke. Front. Hum. Neurosci. 8, 744 (2014).

51. Bizzi, E. & Cheung, V. C. K. The neural origin of muscle synergies. Front. Comput. Neurosci. 7, 51 (2013).

52. Wright, Z. A., Rymer, W. Z. & Slutzky, M. W. Reducing abnormal muscle coactivation after stroke using a myoelectric-computer interface: A pilot study. Neurorehabil. Neural Repair 28, 443–451 (2014).

53. Khorasani, A. et al. Myoelectric interface for neurorehabilitation conditioning to reduce abnormal leg co-activation after stroke: a pilot study. J. Neuroeng. Rehabil. 21, 11 (2024).

54. Sarasola-Sanz, A. et al. Real-Time Control of a Multi-degrees-of-Freedom Mirror Myoelectric Interface During Functional Task Training. Front. Neurosci. 16, 764936 (2022).

55. Daly, J. J. et al. Long-Dose Intensive Therapy Is Necessary for Strong, Clinically Significant, Upper Limb Functional Gains and Retained Gains in Severe/Moderate Chronic Stroke. Neurorehabil. Neural Repair 33, 523–537 (2019).

56. Rodgers, H. et al. Robot assisted training for the upper limb after stroke (RATULS): a multicentre randomised controlled trial. Lancet 6736, 1–12 (2019).

57. Carmena, J. M. Advances in Neuroprosthetic Learning and Control. PLoS Biol. 11, 1–4 (2013).

